# Differential patterns of taxonomic and functional diversity for two groups of canopy arthropods across spatial scales

**DOI:** 10.1101/2022.08.03.502641

**Authors:** Michael B. Mahon, Hannah J. Penn, Kaitlin U. Campbell, Thomas O. Crist

## Abstract

**Aim:** Arthropod diversity is often linked to variation in resource use, dispersal ability, habitat connectivity, and climate factors that differ across spatial scales. The aim of this research was to examine how species richness, functional diversity, and community composition of two taxa differing in functional roles and dispersal ability are structured across spatial scales and to identify the importance of vegetation, climate, and landscape in explaining these patterns at different scales.

**Location:** 96 trees in 24 stands of 6 deciduous forest sites in 2 ecoregions of the eastern USA (North-Central Till Plain and Western Allegheny Plateau)

**Time period:** 2000

**Major taxa studied:** Canopy dwelling ants (Hymenoptera: Formicidae) and spiders (Araneae)

**Methods:** Organisms were collected from tree canopies using insecticidal fogging. Ant and spider taxonomic and functional beta diversity were partitioned across four hierarchical spatial scales (individual tree, forest stand, site, and ecoregion). The contribution of climactic, landscape, and vegetation variables was determined using model selection.

**Results:** Ant and spider species richness, functional diversity, and community composition differed between taxa and across spatial scales. Alpha diversity (within trees) was lower than expected for both taxa and types of diversity, with host tree species supporting different species of ants and spiders. While beta components of species diversity among trees and forest stands was greater than expected for both taxa, spiders also showed significant levels of beta diversity among sites. Functional beta diversity was less scale-dependent than taxonomic beta diversity. Stand-level patterns of beta diversity were significantly predicted by variation in climate and landscape connectivity.

**Main conclusions:** Effects of climate and landscape fragmentation on the diversity and community structure of both taxa indicate that anthropogenic climate change and land use change will alter canopy arthropod communities. Results also suggest that patterns of diversity among fragmentation metrics is influenced by differences in dispersal ability.

## 1 INTRODUCTION

Understanding drivers of species diversity and distributions are primary goals of ecology and can be used to inform conservation efforts. Measures of biodiversity span local (alpha diversity, α) and regional (gamma diversity, γ) scales. Studies of the variation, or turnover, in biodiversity among sites or communities (beta diversity, β), first proposed by Whittaker (1960), have exploded over the last few decades (Anderson *et al*., 2011; Socolar *et al*., 2016; Mammola *et al*., 2021). Beta diversity has been applied to the spatial scaling of biodiversity, temporal change in communities for conservation monitoring, changes across latitudinal and elevational gradients, and environmental filtering of functional traits and subsequent assembly processes (Crist *et al.*, 2003; Kraft *et al*., 2011; Barton *et al*., 2013; Siefert *et al*., 2013; Jarzyna & Jetz, 2018).

Beta diversity has been primarily focused on measures of taxonomic diversity but not functional diversity (i.e. based on the ecological roles and traits of species). To a certain extent, species diversity and functional diversity are correlated, but not reliably so, as not all species are equivalent in a given system and their relative roles may change depending on the presence of and interactions with other species. Species and functional diversity are, therefore, not interchangeable. Indeed, one could imagine two communities comprised of entirely different species (high species beta diversity) that carry out the same functions (low functional beta diversity; Swenson, 2011; Swenson *et al*., 2012; Siefert *et al*., 2013). Because functional diversity quantifies ecological impacts and species interactions, studies including the functional diversity of organisms provide more robust data for conservation and restoration than studies on species presence and abundance alone (Cadotte *et al*., 2009).

Conservation and restoration efforts are landscape-level problems requiring an understanding of how both species and functional diversity scales from local to regional perspectives. Using taxa, such as arthropods, that are widespread, abundant, speciose, and functionally diverse can help researchers test diversity hypotheses across multiple scales (Kemp *et al*., 2017; McCreadie & Adler, 2018). Ants and spiders play several important roles in ecosystem functioning through direct and indirect species interactions (Pearce & Venier, 2006; Maleque *et al*., 2009). Specifically, ants are numerically dominant in most ecosystems, affecting ecological processes through their nest building, predation, and mutualisms (Folgarait, 1998; Crist, 2009); therefore, shifts in ant communities can have multifaceted impacts across trophic levels. Similarly, spiders are obligate predators of other arthropods and may be more susceptible to changes in landscape fragmentation, and loss of key predators is linked to trophic cascades (Bonte *et al*., 2004; Voigt *et al*., 2007).

The mechanisms that might structure arthropod taxonomic and functional beta diversity across spatial scales (e.g., region, site, and forest stand) include characteristics such as land use and topography, management history, and habitat characteristics, respectively. Forest management practices (e.g., logging, fire regimes, and patch size) affect arthropods across multiple scales. At landscape and regional scales, forest fragmentation due to logging or land-use changes lead to decreased habitat area and isolation as well as greater edge to area ratio and altered biotic and abiotic conditions along edge boundaries (Gluck & Rempel, 1996; Fahrig, 2003; Haddad *et al*., 2015). At the stand level, decreased forest area and increased edge cause shifts in tree stand pattern, vegetation structure, and soil characteristics (e.g. moisture, organic matter, texture; Johnston & Elliott, 1996). The effects of these changes on arthropods are many and varied. Fragment size impacts both taxonomic and functional diversity of ants and spiders as well as shifting community composition of these taxa (Pearce *et al*., 2005). Forest habitats with greater tree diversity and structural variability and complexity give rise to a greater variety of niches, and, thus, species and roles to inhabit those niches (Tews *et al*., 2004). Thus, vegetation or habitat complexity can drive patterns of arthropod species richness through direct effects on foraging success and resource availability and indirect effects on ant nest site selection via soil moisture and shading (Mcnett & Rypstra, 2000; Wang *et al*., 2001; Lassau & Hochuli, 2004).

Taken together, beta diversity and composition of key arthropod communities may be affected at a range of spatial scales, potentially driven by several biotic and abiotic characteristics. However, many studies of arthropod diversity focus on a single spatial scale and on taxonomic diversity alone. In this study, we evaluated the role of spatial scale on the taxonomic and functional diversity and the community composition of two arthropod groups (ants and spiders). We quantified these measures across four spatial scales: trees, forest stands, sites, and ecoregions. Specifically, we asked: 1) What spatial scales explain the greatest variation in taxonomic and functional beta diversity and in the community composition of these two taxa? and 2) What are potential mechanisms of scale-associated community structuring, including variation in tree species, forest area and connectivity, and seasonality in climate? We hypothesized that both taxonomic and functional diversity have greater variation at finer spatial scales (tree and stand levels) than at broader spatial scales (site and ecoregion). We also expected that spiders, with greater dispersal abilities, would exhibit greater levels of beta diversity at broader spatial scales than ants, which are more dispersal limited. Similarly, we hypothesized that the biotic and abiotic factors structuring these patterns of diversity differ for spiders and ants given inherent differences in resource use and dispersal ability.

## 2 METHODS

### 2.1 Sampling Design, Study Sites, and Arthropod Collection

Samples were collected from tree canopies of southern Ohio and southeastern Indiana using a hierarchically nested design with four levels. The broadest level – ecoregion – was represented by the glaciated North-Central Till Plain and unglaciated Western Allegheny Plateau (Figure S1). Each ecoregion varies in soil type, forest composition, and topography. The second level – site – comprised 3 sites in each ecoregion for a total of 6 sites (Figure S1). The third level – stand – comprised 4 forest stands within each site (24 total). Within each site, two stands were located in uplands, and two stands were located in lowland topographic positions. The fourth level – tree – included 8 individual trees (96 total) within a 1-ha area of each stand. Each tree was sampled using canopy fogging and arthropods were collected in an array of 12 1-m^2^ funnel arrays placed under the crown of the fogged tree. Arthropods were knocked down by insecticidal fogging (0.5 L of 0.5% pyrethrin-based insecticide) for 3 minutes using a Curtis Dyna-Fogger hoisted into the tree crown and collected by funnels with attached vials of ethanol. Insecticidal fogging is not dependent on arthropod activity and lethality is non-specific (Basset *et al*., 1997; Stork & Hammond, 1997). Sampling was completed during early (22 May-20 June 2000) and late (2-25 August 2000) summer to capture seasonal variation, with early and late season samples pooled for analyses. For more details on sampling methods, see Gering & Crist (2002). Ants were identified to species using the *Ants of Ohio* (Coovert, 2005) and ant functional groups were trait-based (see below) rather than explicitly based on taxonomy or behavior (Andersen, 1997; Crist, 2009). Spider adults were identified to species based on the *Spiders of North America* (American Arachnological Society, 2005), Spiders of Connecticut (Kaston, 1981), and a provisional list of Ohio spiders (Bradley, 2017), while juveniles were identified to family level when possible.

Spider families were used as an indicator of functional guild, based on foraging strategy, prey range, habitat stratification, and circadian activity, as these traits are highly correlated with family (Cardoso *et al*., 2011).

We refer to these samples as canopy or arboreal arthropods because they were collected by fogging tree crowns. Although some of the common species we recorded are primarily arboreal (e.g., *Aphaenogaster mariae,* Table S3), most of the species of ants and spiders are known to move between strata and nest or overwinter in the ground. Nonetheless, Crist and Campbell (2017) recorded significant differences in the community composition of ants from canopy fogging and pitfall traps samples at the same study sites in the North-Central Till Plain.

### 2.2 Data Analyses

#### 2.2.1 Ant functional groups

To classify ants according to functional traits, we selected 10 binary, categorical, and continuous traits that may influence the ecological role of ants (Table S1; (Del Toro *et al*., 2015; Record *et al*., 2018; Mahon, 2019). We used trait definitions and data from Del Toro *et al.* (2015), Record *et al.* (2018), Coovert (2005), and AntWiki (2022). Missing morphological data were supplemented with measurements taken from 2-10 mounted specimens per species collected during this study. Functional groups were formed from a dendrogram based on functional dispersion and the Ward clustering method using the dbFD function (FD package, R; Laliberté & Legendre, 2010); functional groups were delineated by setting six functional groups, where we noted a clear break of functional groups (Table S1, Figure S2).

#### 2.2.2. General patterns and species accumulation curves

For all statistical analyses, early and late sampling periods were pooled and all univariate analyses were conducted in R v4.0.0 (R Core Team, 2022). We constructed sample-based rarefaction curves of ants and spiders by tree species to assess differences in species richness among the most common host trees. To determine the effectiveness of our overall sampling effort, we also conducted species rarefaction and extrapolation curves for ants and spiders using the iNEXT package in R (Chao *et al*., 2014). We also estimated species richness by host trees and overall richness using the Chao1 estimator (Chao, 1984; Colwell & Coddington, 1994).

#### 2.2.3 Diversity partitioning

To analyze taxonomic diversity, we partitioned species richness and functional group richness (q = 0) across hierarchical levels, with multiplicative partitioning methods based on the PARTITION software developed by Crist *et al.* (2003) using the R package, PARTITIONR (Mahon *et al*., 2019). Multiplicative partitions express beta diversity as the effective number of distinct communities, whereas alpha diversity is in units of species richness (Anderson *et al*., 2011). Using hierarchical diversity partitioning, beta components can be separated into nested hierarchical levels (i.e. spatial scale). Hierarchical multiplicative partitioning calculates beta diversity at a given level (i) by dividing the average alpha at the i+1 level, *β_i_* = *α*_*i*+1_/*α_i_*. We tested the significance of *α*_1_ (within trees), *β*_1_ (among trees), *β*_2_ (among stands), *β*_3_ (among sites), and *β*_4_ (between ecoregions) against null, random distributions using 1000 sample-based randomizations. This type of randomization preserves intraspecific aggregation at each hierarchical level, and thus tests the null hypothesis that observed patterns of species diversity at each level are similar to those expected by randomized aggregation of samples at each level of the hierarchical design; alternatively, the null hypothesis is rejected if observed patterns are significantly different from those expected from null distributions, supporting non-random hierarchical species assemblages that are structured by ecological processes (Crist *et al*., 2003). Partitioning of alpha, beta diversity in this manner therefore accounts for issues of spatial pseudoreplication that can arise in local-to-regional comparisons of species richness (Gering & Crist, 2002). We used a two-tail probability of *p* = 0.05 *(p* = 0.025 for each tail of the null distribution) to determine whether observed diversity patterns were higher or lower than expected via randomizations. To compare relative deviations of the null distributions from the expected values across hierarchical levels and endpoints, we calculated standard effect sizes (SES; also termed beta and alpha deviations) from the mean and standard deviation of the null distributions for each hierarchical level (*SES* = (*I_obs_* — *I_exp_*)/*σ*) (Gotelli & McCabe, 2002; Kraft *et al*., 2011).

To analyze variation in community composition across hierarchical levels, we conducted analyses for multivariate location using PRIMER-E and PERMANOVA+ v6 (Anderson, 2001). We used permutational multivariate analysis of variance (PERMANOVA) to partition the variation in community composition across hierarchical levels. PERMANOVAs were conducted using 9999 permutations of the data, with a nested design. Since multivariate analyses were conducted on square-root transformed abundance data with a Bray-Curtis dissimilarity, variance components of PERMANOVA can be interpreted as percent dissimilarity (Anderson, 2001).

#### 2.2.4 Landscape and environmental variable analyses

To address potential mechanisms of diversity structuring, we compiled and evaluated landscape, climactic, and vegetation variables for forest stands and surrounding areas. Landscape variables were habitat fragmentation measures (estimated using ESRI arcGIS Dekstop v10.6.1 (ESRI, 2019), Fragstats v4.2.1 (McGarigal *et al*., 2012)) and land use data (USDA NASS Crop data layer for 2008, the closest year with most reliable data for both Indiana and Ohio (USDA National Agricultural Statistics Service, 2022)). Land cover/land use composition and configuration was collected for of 1.0, 3.0, 6.5, and 10.0 km buffers surrounding each stand to accommodate the potential dispersal distances of both spiders and ants, and to minimize overlap between stand buffers (Thomas *et al*., 2003). Initial model selection (see methods below) was conducted to determine which buffer size explained the most variation for both ants and spiders; data at the 6.5 km buffer were used for the remainder of the statistical analyses. Each landscape was analyzed using a “no-sampling” strategy and an 8-cell neighborhood rule for the following class level metrics as specified in Fragstats: CLUMPY, PLAND, GYRATE_MN, GYRATE_AM, and PARA (McGarigal *et al*., 2012). CLUMPY (fragmentation index) is an index of fragmentation of deciduous forest within the measured landscape where −1 is highly fragmented forest and 1 is a complete, unfragmented forest; the CLUMPY index is not confounded by changes in forest area. PLAND is the percentage of deciduous forest within the landscape. GYRATE_MN is the mean distance (m) to forest edge from patch centroid. GYRATE_AM (patch connectedness), is the area-weighted distance (m) to forest edge from patch centroid, or patch connectedness. PARA (patch edge:area ratio) is the mean edge:area ratio for all deciduous forest patches. We obtained 19 bioclimatic variables from WorldClim v2.0 (Fick & Hijmans, 2017); due to the relatively course resolution (~1 km^2^) of the bioclimatic variables, 2 of our stands (within Brookville) had identical climatic data, but all other stands and sites varied. We included a stand-level vegetation measure of tree species richness (dbh ≥10 cm) by recording all tree species present within the same 1-ha stands where trees were fogged (Crist *unpublished data).*

We accounted for collinearity among landscape and environmental variables by removing those that were highly correlated (Pearson *r* ≥ 0.80). In total, 10 variables were included: 4 landscape variables, 5 climatic variables, and 1 vegetation variable (Table S2). Prior to all analyses, variables were standardized to z-scores to aid in model fitting and inference. To determine climatic, landscape, and vegetative influence on patterns of taxonomic and functional richness, we used linear regression models (lm function, stats package, R; (R Core Team, 2022) with response variables of mean alpha (*α*_1_, within trees) and beta (*β*_1_, among trees) diversity for taxonomic and functional diversity at each stand. Model selection allowed for the additive term of all environmental predictors. To identify best models, we used the lowest Akaike’s Information Criterion with bias-correction (AICc). For best models, we calculated AICc weights (*w*) and appropriate R^2^. We tested for spatial autocorrelation using Moran’s I (Moran.I function, ape package, (Paradis & Schliep, 2019), which indicated no spatial autocorrelation present in the residuals of our univariate models.

Similarly, we used DISTLM and distance-based redundancy analyses (dbRDA) in PRIMER and PERMANOVA+ (McArdle & Anderson, 2001) to assess influences of our environmental variables on community composition at the stand level. We performed stepwise model selection based on AIC values to examine the relationship between the explanatory variables and community composition. We only included variables with significance of *p* < 0.10 in preliminary marginal tests to reduce likelihood of overfitting multivariate analyses.

Significance values and variance explained by selected predictors were found using bootstrap tests based on 9999 iterations. Multivariate analyses were conducted on square-root transformed abundance data with Bray-Curtis dissimilarity.

## 3 RESULTS

We collected 3,053 individual ants representing 23 species with 2 singleton species (Table S3). Estimated Chao1 richness was 24 ± 2 species, with rarefaction curves indicating ant richness had plateaued (Figure S3). We collected 5,221 individual spiders representing 23 families, 67 genera, and 97 species. Of these, 83.7% were juveniles, we identified the remaining 925 adults to morphospecies (Table S3). Of these species, 15 were represented by a single specimen. Dominant spider families were Araneidae (23%), Linyphiidae (17%), Salticidae (16%), Anyphaenidae (12%), and Theridiidae (10%). Estimated Chao1 richness was 108 ± 6 species, with rarefaction indicating spider richness began to plateau and likely would have been reached by 150 tree samples (Figure S3).

### 3.1 Hierarchical diversity partitioning

Spider species alpha and beta diversity were higher than ant species diversity, ant functional diversity, and spider functional diversity (Table 1). Multiplicative beta diversity based on species richness (q = 0) exhibited similar patterns between taxa, with beta decreasing as scale increased (Figure 1). For species richness, the observed β_tree_ components were 2x the α_tree_ of 4.8 species of ants per tree and 2.8x of 6.4 species of spiders per tree, respectively. The β_stand_ values were 1.6x the combined of 9.5 species of ants per stand, and 2.4x the 17.9 species of spiders per stand. The β_site_ components were 1.3x the 15.7 species of ants per site and 1.7x the 42.9 spider species per site, and β_ecoregion_ were <1.3x the mean richness of ants and spiders per ecoregion (Table 1). The functional group diversity components mirrored those of species richness, except that the α_tree_ and β_tree_ levels comprised most of variation in total functional diversity of ants with additional variation explained by β_stand_ for spiders. Randomization tests of species richness indicated α_tree_ was significantly lower than expected across taxa (*p* < 0.001), β_tree_ was higher than expected across taxa (*p* < 0.001), β_stand_ was higher than expected for ant species and spiders (*p* < 0.001), β_site_ was higher for spider species (*p* = 0.005), and β_region_ was not different across taxa (*p* > 0.025, Table 1). These patterns emerged, despite having similar diversity across scales for diversity for ant species, ant functional groups, and spider functional groups (Figure 1, Table 1). At local scales (within and among trees), for both taxonomic and functional endpoints, the deviations from observed diversity were greater for ants than spiders, but the opposite pattern emerged at broader scales (stand, site, and region; Figure 1), indicating differential hierarchical patterns of species diversity between ants and spiders.

**Figure 1.**
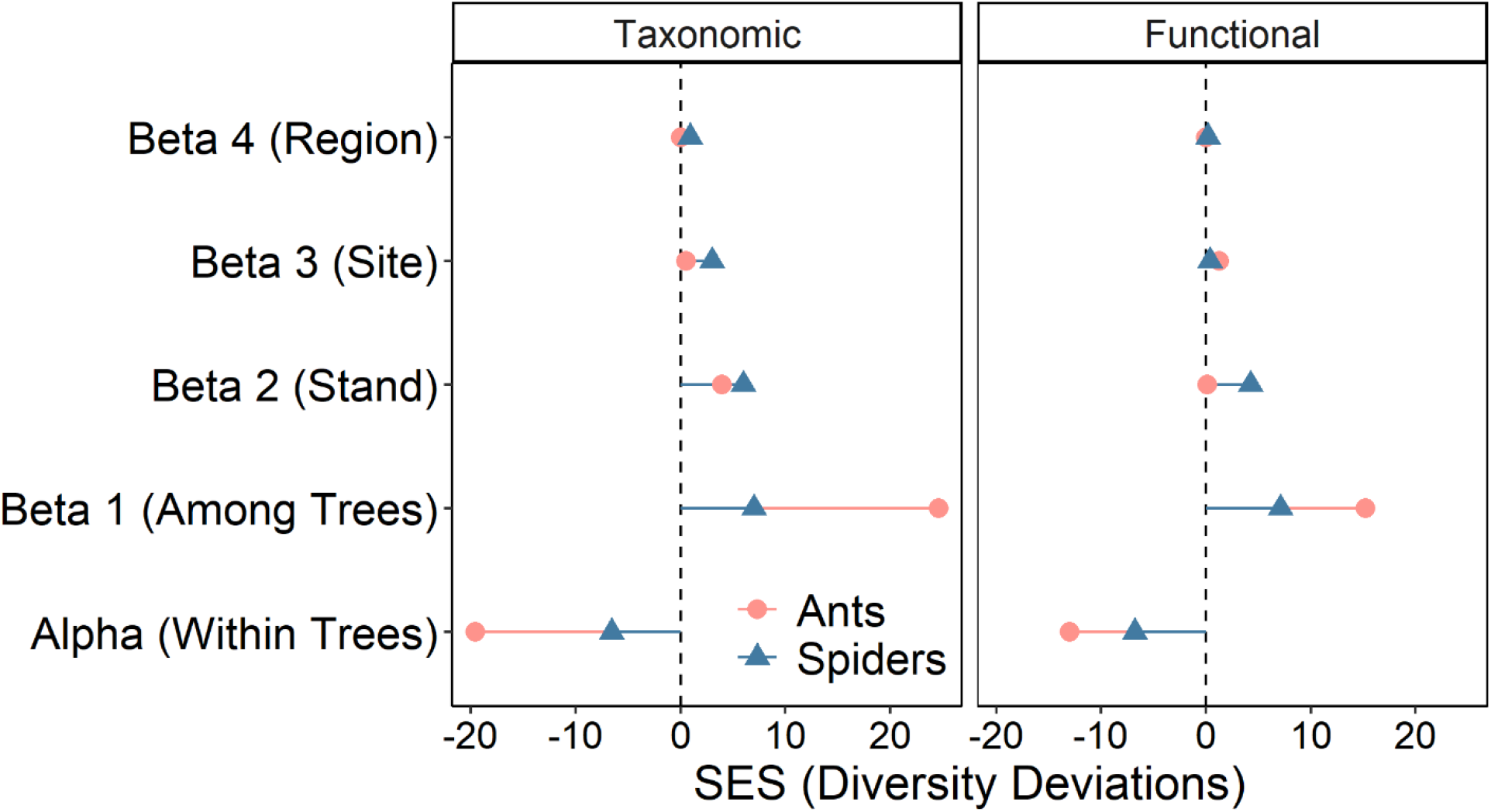
Standard effect sizes (SES) of the observed diversity partitions and the diversity deviations from a null model across the hierarchical scales (within trees, among trees, stand, site, and region) for taxonomic and functional diversity. Blue triangles are spiders, pink circles are ants.

**Table 1.**
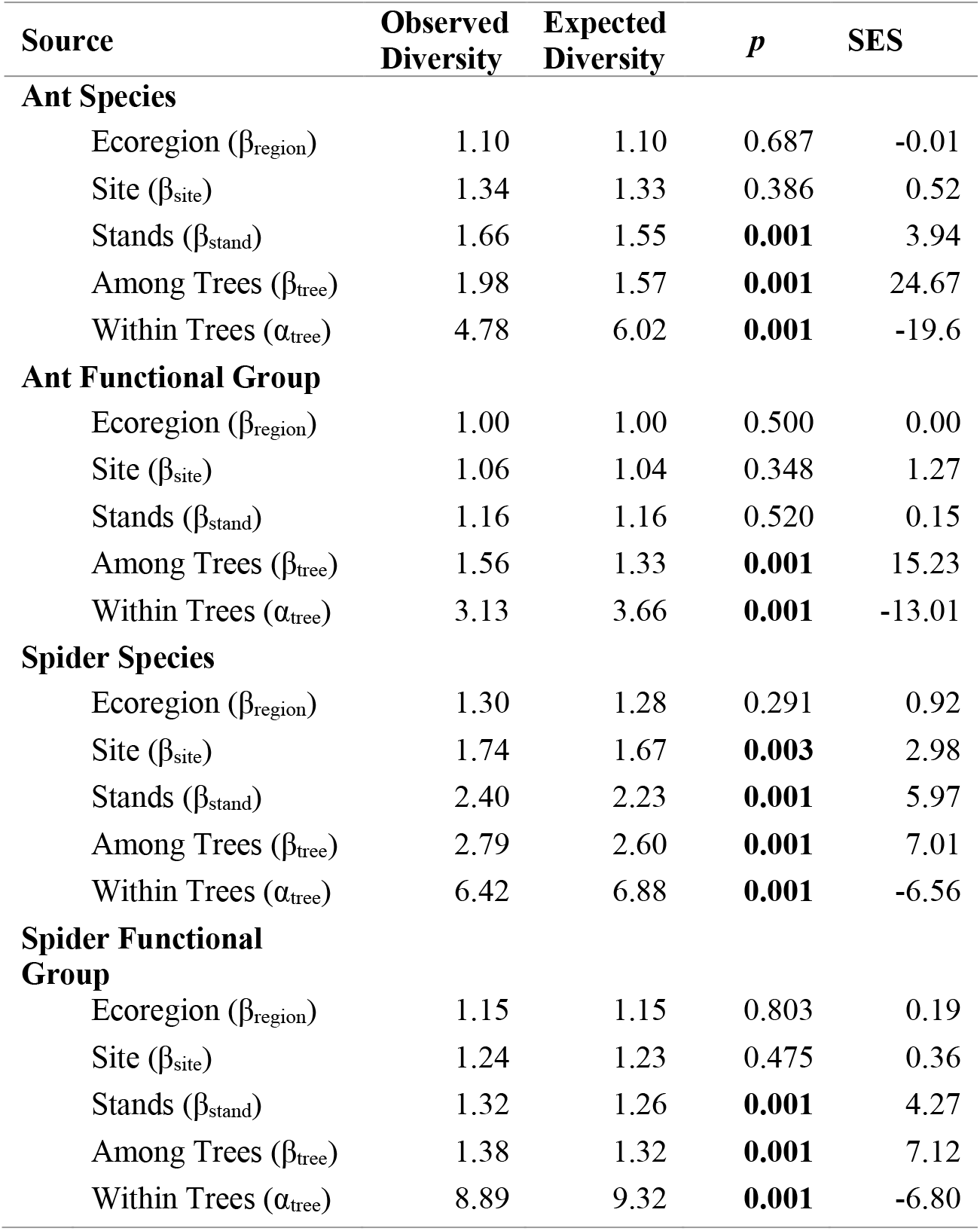
Multiplicative partition of alpha and beta diversity components across a hierarchically nested study of canopy ant and spider diversity. *P* values <0.05 are bolded

### 3.2 Community composition

We tested for multivariate differences in ant and spider species composition across hierarchical levels based on variation in Bray-Curtis dissimilarities (Table 2). For both taxa, a greater proportion of the variation in dissimilarities was explained at site and ecoregion levels than with univariate diversity partitions, but components still decreased with increasing spatial scale (Table 2). Spiders also showed greater residual variability among trees (56%) compared to ants (40%). Similarly, spider species composition showed greater variance among stands within sites than ant species composition (Table 2). Ant assemblages showed significant variability across all spatial scales; however, this was not the case for spiders, as the variance component for ecoregion was not different from zero (Table 2).

**Table 2.** Permutational multivariate analyses of variance (PERMANOVA) based on Bray-Curtis dissimilarity of square-root transformed abundance data for species and functional groups for both taxa. *P*-values are based on Monte-Carlo randomization, *p* values <0.05 are bolded. Variance is the square root of the estimated component of variance, to put values on the scale of Bray-Curtis dissimilarities (percent difference among assemblages).

| Endpoint and Source of Variation | *df* | MS | Pseudo-F | *p* | Variance | Percent Variance Explained |
| --- | --- | --- | --- | --- | --- | --- |
| **Ant Species** |  |  |  |  |  |  |
| Ecoregion | 1 | 14407 | 2.41 | **0.042** | 13.3 | 15.9% |
| Site(Ecoregion) | 4 | 5970.3 | 2.31 | **<0.001** | 14.5 | 17.4% |
| Stand(Site(Ecoregion)) | 18 | 2589.7 | 1.65 | **<0.001** | 16.0 | 19.2% |
| Residual | 72 | 1566.1 |  |  | 39.6 | 47.5% |
| **Ant Functional Groups** |  |  |  |  |  |  |
| Ecoregion | 1 | 10354 | 2.71 | 0.066 | 11.7 | 17.8% |
| Site(Ecoregion) | 4 | 3821.6 | 2.61 | **0.003** | 12.1 | 18.5% |
| Stand(Site(Ecoregion)) | 18 | 1463.4 | 1.63 | **0.003** | 11.9 | 18.1% |
| Residual | 72 | 899.2 |  |  | 30.0 | 45.7% |
| **Spider Species** |  |  |  |  |  |  |
| Ecoregion | 1 | 15822 | 1.82 | 0.058 | 12.2 | 11.7% |
| Site(Ecoregion) | 4 | 8706 | 1.82 | **<0.001** | 15.7 | 15.1% |
| Stand(Site(Ecoregion)) | 18 | 4778.7 | 1.53 | **<0.001** | 20.4 | 19.6% |
| Residual | 72 | 3120 |  |  | 55.9 | 53.7% |
| **Spider Functional Groups** |  |  |  |  |  |  |
| Ecoregion | 1 | 4849.3 | 2.01 | 0.089 | 7.1 | 13.5% |
| Site(Ecoregion) | 4 | 2415.2 | 1.80 | **0.023** | 8.2 | 15.5% |
| Stand(Site(Ecoregion)) | 18 | 1343.8 | 2.42 | **<0.001** | 14.0 | 26.5% |
| Residual | 72 | 554.9 |  |  | 23.6 | 44.5% |

For both ants and spiders, functional composition was less variable than taxonomic composition across scales (Table 2). Ant and spider functional assemblages were significantly variable at the stand and site levels, but the variance component for ecoregion was not different from zero (Table 2). While spider functional composition showed reduced variance components with increasing spatial scale, ant functional composition showed little differentiation among stand, site, and ecoregion scales, but variance was highest among sites (Table 2), indicating stronger variability across sites within ecoregion than among stands within sites. Nonetheless, both ant and spider functional composition was most variable among trees within stands (Table 2), suggesting functional differentiation of these communities among tree species.

### 3.3 Patterns of diversity and community composition

Tree identity influenced spider and ant species richness. For ants, white oak *(Quercus alba*), red oak (*Q. rubra*), American beech (*Fagus grandifolia*), sugar maple (*Acer saccharum*), and hickory *(Carya* spp.) hosted similar levels of richness, while tulip poplar *(Liriodendron tulipifera*) maintained the lowest richness (Figure 2B). Ant richness began to plateau on most tree species (but see hickory, Figure 2A). For spiders, white oak, sugar maple, and American beech had greater richness than tulip poplar, red oak, and hickory, as indicated by rarefaction (Figure 2B). Unlike for ants, most tree species did not begin to plateau for spiders, except for red oak.

**Figure 2.**
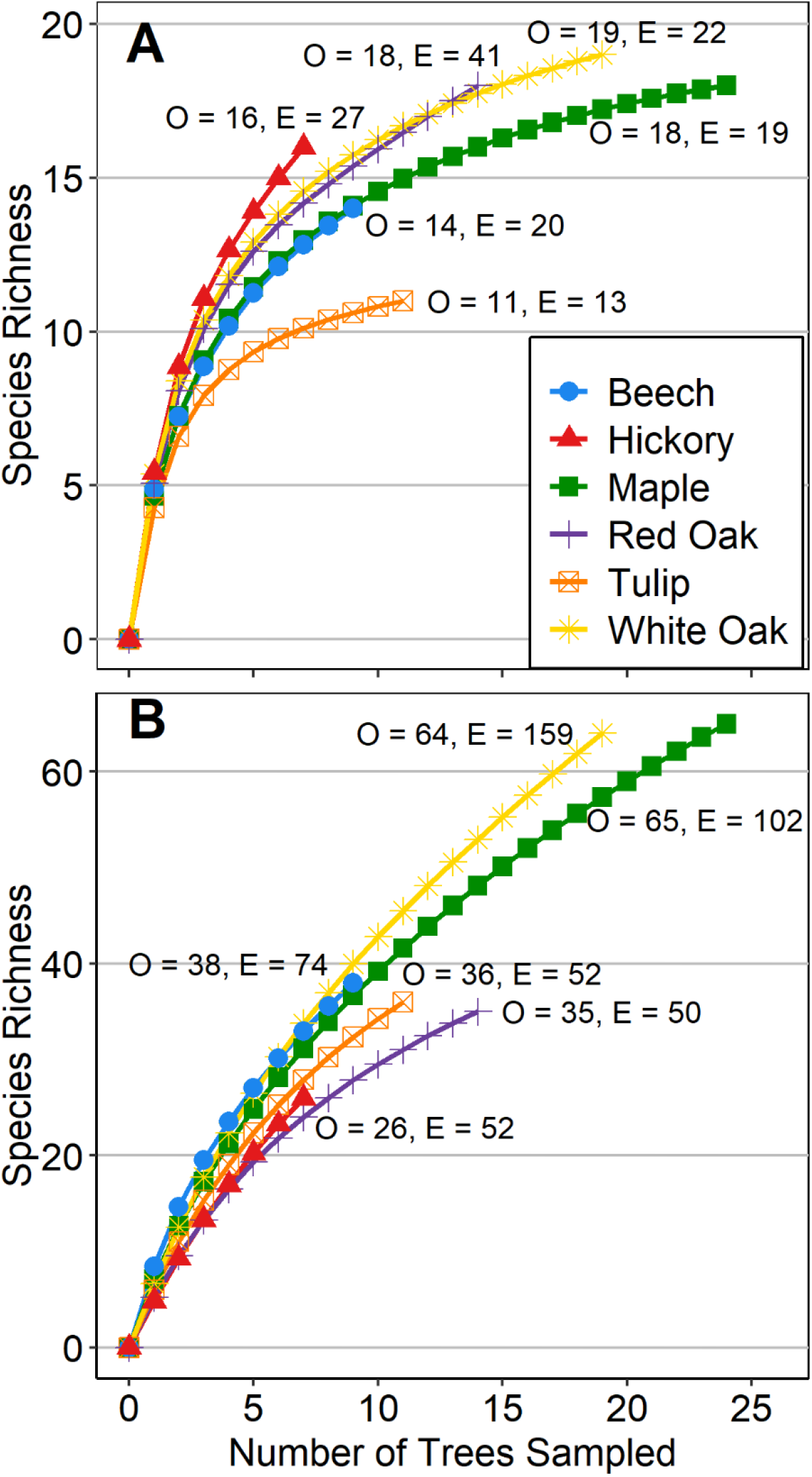
Sample-based rarefaction curves of ant (A) and spider (B) species by number of host trees sampled. Ash, Hackberry, Sycamore, and Walnut trees are not shown, because of few individual trees sampled (< 5 for each species). Number of points on a curve and length of the curve represent the number of individual trees fogged. Ants and spiders show differential patterns among tree species of observed (O) and Chao1 estimated (E) species richness.

Average alpha (α_1_, within trees) and beta (β_1_, among trees within stand) diversity for taxonomic and functional endpoints were driven by a combination of landscape and climatic variables measured at the stand level. Temperature evenness (isothermality) and precipitation seasonality were important predictors of alpha diversity of spider functional groups, ant species, and ant functional groups as well as beta diversity of spider species, spider functional groups, and ant species (Table 3). Specifically, precipitation seasonality was negatively associated with alpha diversity, but positively associated with beta diversity (Table 3). This suggests that increased variability in precipitation may simultaneously reduce average alpha within trees while increasing differentiation of arthropod communities among tree species (beta). Stand-level richness of tree species was positively associated with beta diversity of ant species, but negatively associated with beta diversity of spider functional groups (Table 3). Finally, patch connectedness, size, and shape were important predictors of alpha diversity of ant and spider functional groups and of beta diversity of ant functional groups (Table 3).

**Table 3.** Multiple regressions of tree-level alpha and beta diversity as predicted by stand-level measurements of climate, landscape structure and tree species richness. Regression coefficients of best models as indicated by AICc model selection. All predictor variables were converted to Z scores (SE) to allow for comparison of strength of effects.

|  | Predictor Variables |  |  |  |  |  |  |  |
| --- | --- | --- | --- | --- | --- | --- | --- | --- |
|  | df | Patch<br>Connectedness | Patch<br>Edge:Area | Tree<br>Richness | Isothermality | Temp Warm<br>Month | Precipitation<br>Seasonality | R<sup>2</sup> |
| Alpha |  |  |  |  |  |  |  |  |
| Ant Species | 2 |  |  |  |  |  | -0.73 (0.22) | 0.53 |
| Ant Func. Group | 3 | 0.44 (0.18) |  |  |  |  | -0.60 (0.18) | 0.72 |
| Spider Species | 1 |  |  |  |  |  |  | --- |
| Spider Func. Guild | 3 |  | -0.99 (0.25) |  |  |  | -1.27 (0.24) | 0.70 |
| Beta |  |  |  |  |  |  |  |  |
| Ant Species | 4 | 0.82 (0.21) |  | 0.49 (0.20) |  |  | 0.86 (0.21) | 0.63 |
| Ant Func. Group | 2 |  |  |  |  | -0.58 (0.26) |  | 0.33 |
| Spider Species | 2 |  |  |  | -0.54 (0.27) |  |  | 0.29 |
| Spider Func. Guild | 3 |  |  | -0.52 (0.23) |  |  | 0.49 (0.23) | 0.44 |

Similar to univariate patterns of alpha and beta diversity, variation in multivariate community composition using dbRDA was driven by a similar combination of landscape and climatic drivers. Variation in ant community composition was best explained by precipitation variability (R^2^ = 16.3%) (Figure 3A). Variation in ant functional group composition was best explained by isothermality (R^2^ = 40%), patch connectedness (14.2%), and precipitation seasonality (11.1%) (Figure 3B). Variation in spider community composition was best explained by isothermality (R^2^ = 17.6%), maximum temperature (15.6%), and habitat fragmentation (10.4%) (Figure 3C). Variation in spider functional guild composition was best explained by patch connectedness (R^2^ = 25.2%), isothermality (16%), stand tree richness (9.6%), and habitat fragmentation (8.9%) (Figure 3D).

**Figure 3.**
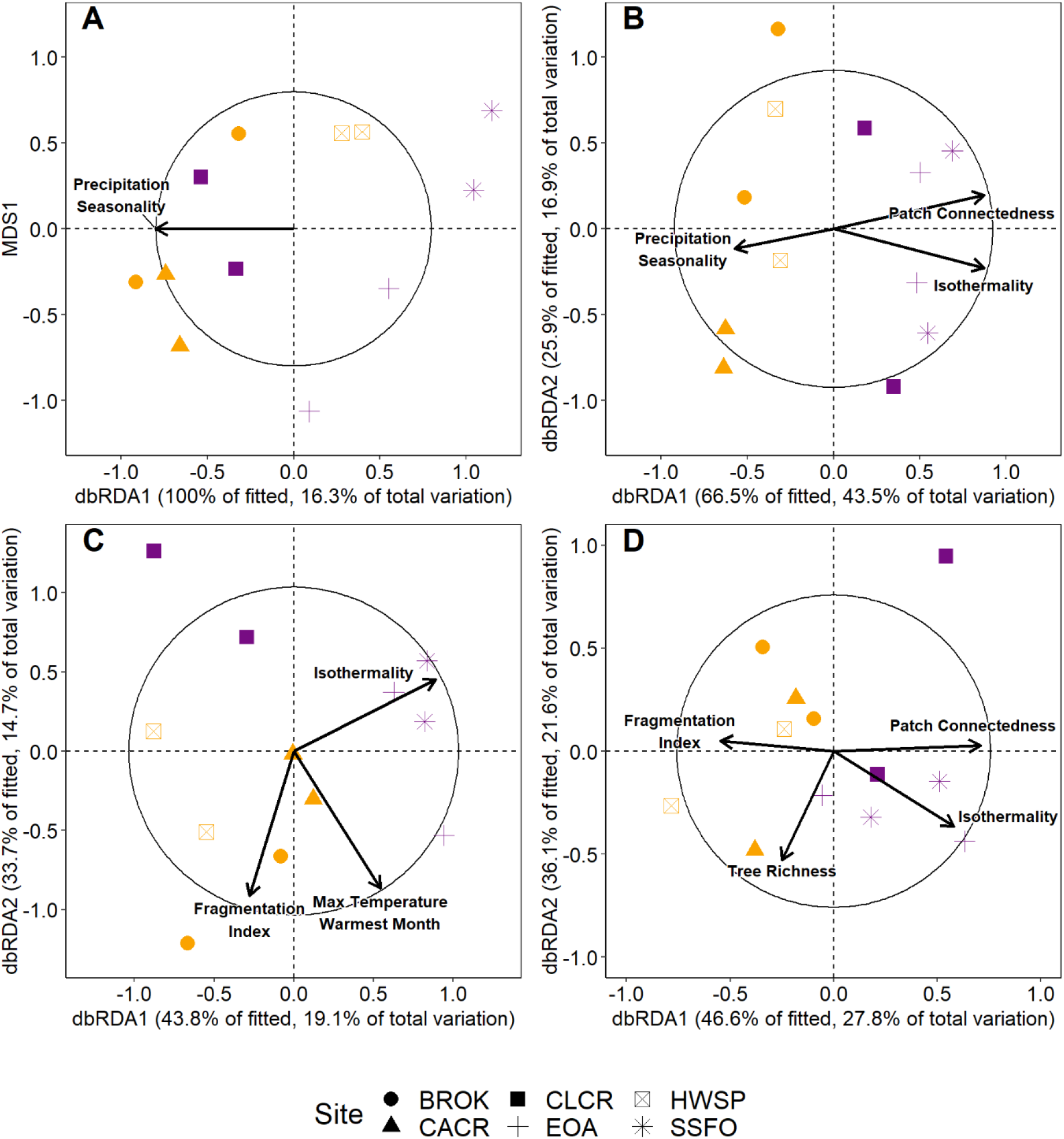
Distance-based redundancy analysis (dbRDA) of stand-level variation in composition based on Bray-Curtis dissimilarities of square-root transformed abundance of ant species (A), ant functional groups (B), spider species (C), and spider functional groups (D). Vectors correspond to predictor variables in best model as determined by model selection. The circles correspond to vector lengths that would have a correlation coefficient of 1 with a given axis, strength of correlation for vectors are scaled to this circle. Yellow symbols are stands within sites in the North-Central Till Plains and purple symbols are stands within sites in the Western Allegheny Plateau. Sites are designated by different symbol shapes.

## 4 DISCUSSION

### 4.1 Spatial structure of diversity

Our results indicate differential patterns of beta diversity and community structure between canopy-dwelling ants and spiders and that taxonomic diversity is more variable at larger spatial scales than functional diversity. In support of our hypothesis that spider diversity would be more affected by broader spatial scales than ants, we saw greater diversity and greater deviation from expected diversity (SES) at broader spatial scales for spiders than for ants, but there was more variation between ecoregions for ant communities than for spider communities. While functional and taxonomic diversity exhibited similar patterns across scales, taxonomic diversity exceeded functional diversity at coarser spatial scales (Site and Ecoregion), suggesting trait clustering and functional redundancy at broader spatial scales, consistent with the findings of Jarzyna & Jetz (2018). Essentially, the loss or addition of species at the site and ecoregion scales has little influence on the loss or addition of ecological functions – as determined by functional groups – at these same scales.

The finding that broad-scale (Ecoregion) beta components of diversity were not different than expected suggests the effects of ecoregions do not structure the taxonomic and functional diversity of canopy-dwelling ants and spiders. Our results contrast with the significant differentiation of beetle diversity between ecoregions from these same samples (Gering *et al*., 2003) but are consistent with those of Summerville *et al.* (2003) who found no significant deviation from expected diversity in forest moth species richness among ecoregions. Nonetheless, both ant and spider communities showed similar patterns of decreasing community variation with increasing spatial scale, indicating strong differences in beta diversity at the tree and stand levels, consistent with the findings of Gering *et al.* (2003) and Summerville *et al.* (2003). Here, we also conducted multivariate partitions of Bray-Curtis dissimilarity across scales, which showed slightly larger components of variation at the site and ecoregion levels than univariate partitions. This suggests that shifts in the relative abundance of species or functional group across sites and ecoregions were more important than shifts in species or functional group composition.

Lower variation than expected at the individual tree-level implies that tree species support distinct levels of species richness and, likely, functional diversity of arthropod taxa, as supported by rarefactions of individual tree species for both ants and spiders. Differences in tree species-arthropod species richness relationships (estimated and observed richness) indicate potential tree-species specific constraints such as prey species abundance and competition/territoriality in ants (Majer & Delabie, 1999; Yasuda & Koike, 2009) and nesting/web site limitations for spiders (Nicolai, 1986; Larrivée & Buddle, 2010). Thus, the maintenance of temperate, canopy-dwelling arthropod communities is, at least somewhat, dependent on maintaining the diversity of host trees. As such, the subsequent emerald ash borer *(Agrilus planipennis)* driven loss of ash trees *(Fraxinus* spp.; Herms & McCullough, 2014) and long-term declines of several oak species in eastern forests (McEwan *et al*., 2011) since the time of sampling has likely resulted in the loss of distinct arboreal arthropod communities and, possibly, species since the early 2000s.

### 4.2 Landscape and environmental influences

Previous studies have found differential environmental drivers between taxonomic and functional diversity (Longhi & Beisner, 2010; Pool *et al*., 2010; Meynard *et al*., 2011), suggesting that different environmental filters act on taxonomic and functional diversity of the same taxa. While we saw similar scaling patterns between ants and spiders, taxonomic and functional diversity, and community structure, the climatic, landscape, and vegetation drivers of these patterns did, indeed, differ (Table 3, Figure 3). Climatic characteristics showed strong influence on alpha and beta diversity as well as both taxonomic and functional composition, while landscape characteristics were important in explaining patterns of functional alpha diversity and functional composition. Nonetheless, taxonomic and functional community composition was explained by shared climatic variables (precipitation seasonality and isothermality), indicating that both taxonomic and functional assemblages are partially driven by the same climatic conditions.

Climate variables were more common than landscape variables in univariate (diversity) and multivariate (community composition) ant and spider models. Precipitation seasonality was the most common predictor in explaining variation in taxonomic and functional diversity (Table 3), while isothermality (temperature evenness) was the most common predictor for explaining multivariate variation in taxonomic and functional community composition (Figure 3). These differences in climate variables as predictors of univariate and multivariate variation in diversity may also reflect variation in species distributions versus shifts in relative abundances. The broader implication of climate variability as an important driver of diversity and composition of canopy arthropod communities is that climate change mediated increases in climate variability (increased precipitation seasonality and decreased isothermality) will likely greatly alter these canopy-dwelling communities (Westerling, 2016; Jump *et al*., 2017; Neumann *et al*., 2017).

Habitat configuration was more important in explaining patterns of ant and spider alpha and beta diversity and community structure than total habitat availability, but spider functional richness was related to patch size and shape. In line with our results, previous research suggests that ants show weak support for species-area relationships, while spiders tend to exhibit strong positive species-area relationships (Crist, 2009; Cardoso *et al*., 2010). Similarly, previous studies have found higher ant richness in habitat fragments with higher connectivity, while habitat connectivity does not influence spider species richness (Abensperg-Traun *et al*., 1996; Suarez *et al*., 1998; Cardoso *et al*., 2010). The disparities in the influence of habitat fragmentation and patch size and shape between ants and spiders is likely due to differences in dispersal ability (Thompson & Townsend, 2006). Our results suggest that patterns of arthropod diversity among habitat patches is influenced by dispersal ability, with connectivity being an important predictor of poor dispersing arthropods and patch size and shape being an important predictor of diversity of stronger dispersing arthropods. Taken together, this suggests that the importance of habitat configuration and area of habitat in determining species richness (Haddad *et al*., 2017) may be somewhat dependent upon the dispersal ability of organisms. Yet, connectivity and fragmentation of habitat patches were important for composition of both taxa, suggesting that while spider richness may not be influenced by habitat configuration, the identities of spider species is influenced by habitat configuration. Therefore, conservation efforts should focus on increasing both patch connectivity and size to provide the largest benefits to diversity of all arthropods. Nevertheless, future research should expand upon the relative roles of habitat configuration and patch size and shape in driving patterns of diversity for taxa across a dispersal gradient.

### 4.3 Conclusions

Our findings demonstrate stronger scaling patterns of taxonomic diversity than functional diversity from local to regional scales, suggesting functional redundancy at broader spatial scales. Further, taxonomic and functional diversity and community assemblages change along different environmental gradients. The controls of climate and landscape fragmentation on the diversity and structure of canopy-dwelling ants and spider communities indicate that climate change via increased variability will likely further alter the diversity, composition, and function of arboreal arthropods that are already threatened by forest fragmentation and land use changes. Our findings provide further support for the consideration of functional components of diversity and multiple measures of beta diversity in monitoring and conservation (Cadotte *et al*., 2011; Socolar *et al*., 2016; Isbell *et al*., 2017; Jarzyna & Jetz, 2018).

## Acknowledgements

The Nature Conservancy Ecosystem Research Program, The Ohio Board of Regents Research Challenge Program, and the Miami University Committee on Faculty Research provided funding for this project. Insect sampling was conducted by J. Gering, C. Yeager, N. Anderson, J. Kahn, J. Veech, K. Summerville, and D. Golden. We would also like to thank A. Schaefer, A. VanGorder, K. Donahue, G. Dahlem, and M. Crist for sample processing. Mention of trade names or commercial products in this publication is solely for the purpose of providing specific information and does not imply recommendation or endorsement by the U.S. Department of Agriculture. USDA is an equal opportunity provider and employer.

## Authorship

TOC originally conceived of the study and collected samples; all authors conceived of current manuscript; HJP and KUC identified spiders and ants from samples; MBM conducted statistical analyses; MBM, HJP, and KUC wrote the first draft of the manuscript; all authors revised the manuscript

## Biosketch

This research team aims to better understand and quantify arthropod community structure and diversity at landscape scales through local experiments and broad scale observational approaches. They are interested in refining our understanding of the structuring of biodiversity to improve conservation efforts.

## Code and Data accessibility

All data and R code are available on github: https://github.com/mahonmb/CanopyArthropodDiversity. PRIMER and PERMANOVA+ code is available by request.

## Notes

https://github.com/mahonmb/CanopyArthropodDiversity

### Competing Interest Statement

The authors have declared no competing interest.

