## Supplementary material for "Differential patterns of taxonomic and functional diversity for two groups of canopy arthropods across spatial scales": Supp.

Appendix 1.

Figure S1. Map of Ohio and surrounding states (Indiana, Kentucky, and West Virginia) showing the location and relative distances among the three sampling sites in each ecoregion. Coloration is representative of ecoregions. Solid lines represent state boundaries.

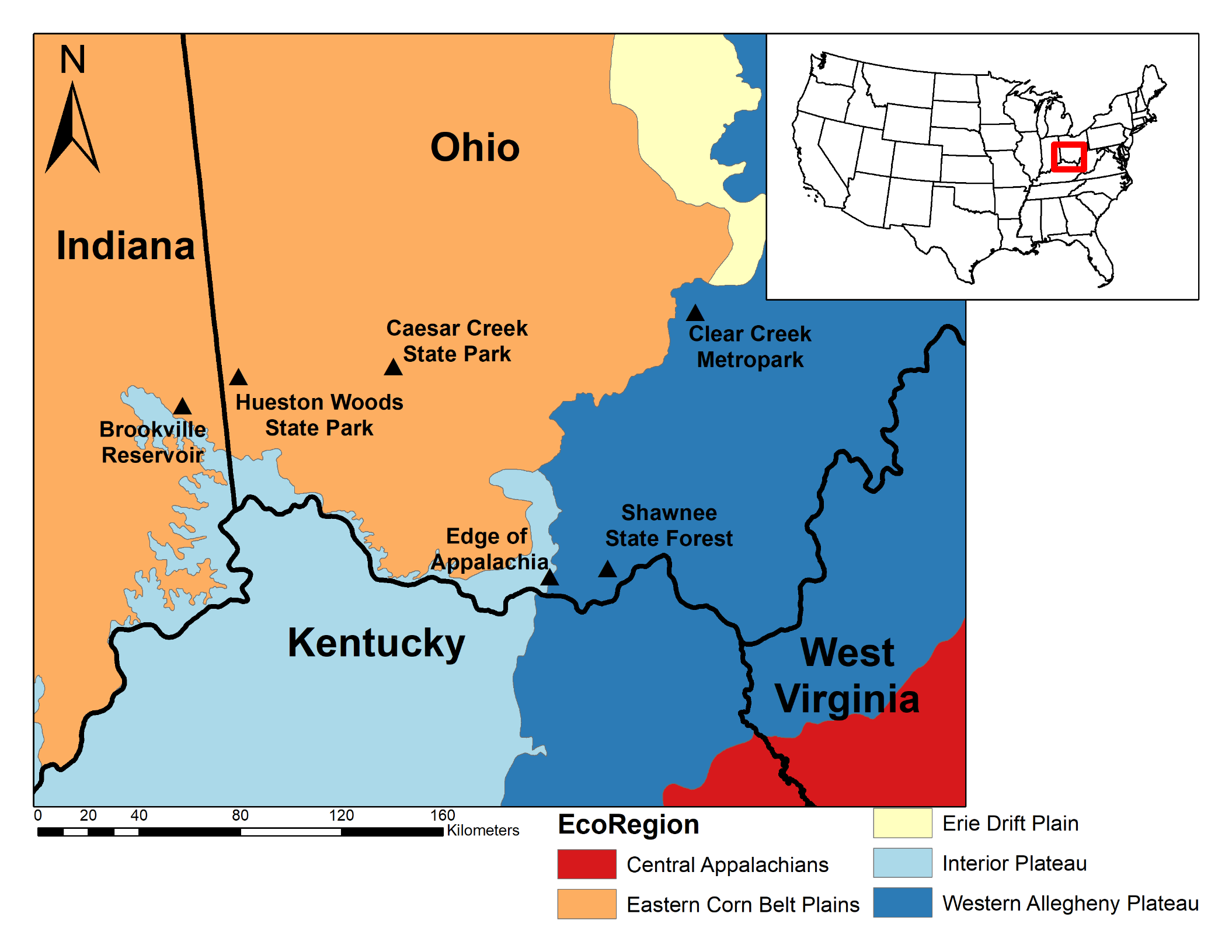

Table S1. Descriptions of traits and functional roles for the 6 ant functional groups found in this study. We clustered the 23 ant species found in this study into functional groups based on morphometric and natural history traits. We used trait definitions and data from Del Toro et al. (2015), Record et al. (2018), Coovert (2005), and AntWiki (2022).

| Functional Group | Primary Functional Role | Characteristics |
| --- | --- | --- |
| Group 1 | Social Parasites | Social parasitic, Medium body size, Forest Dwelling, wood dwelling |
| Group 2 | Seed Dispersers | Medium body size, Forest Dwelling, omnivorous, seed dispersing |
| Group 3 | Soil Movers | Soil dwelling, medium to large colony size, omnivores, |
| Group 4 | Large Cavity Nesters | Large body sized, omnivorous, wood dwelling, forest |
| Group 5 | Small Cavity Nesters | Small body size, acorn and wood dwelling, forest species, small colonies |
| Group 6 | Invertebrate Community Regulators | Omnivores, soil dwelling, forest and edge species, small to medium body size, medium to large nests |

Table S2. Environmental Variables used in dbRDA and linear models. Landscape variable names are those listed in Fragstats with labels used throughout text for ease of interpretation in parentheses (McGarigal *et al.*, 2012).

| Variable | | Explanation | Interpretation |
| --- | --- | --- | --- |
| Landscape | |  |  |
|  | CLUMPY (fragmentation index) | Measure of dispersion of deciduous forest | Range from -1 (maximally disaggregated forest patches) to 1 (entire landscape is forest) |
|  | GYRATE_MN | Mean distance between each cell in the patch and patch centroid and range of these values in the landscape. | Indication of patch extent |
|  | GYRATE_AM  (Patch connectedness) | Area-weight mean patch radius of gyration; measure of landscape connectivity. | Average distance an organism can travel from a random point moving in a random direction |
|  | PARA_MN  (Patch edge:area ratio) | Mean perimeter to area ratio across patches |  |
| Climatic | |  |  |
|  | Isothermality | Quantifies size daily temperatures oscillate relative to annual oscillations. | An isothermal value of 100 indicates the diurnal temperature range is equivalent to the annual temperature range, while anything less than 100 indicates a smaller level of temperature variability within an average month relative to the year. |
|  | Temp Warm Month | Max temperature of warmest month |  |
|  | Temp Dry Quarter | Mean temperature of driest 3 months |  |
|  | Precipitation Wettest Quarter | Total precipitation of wettest 3 months |  |
|  | Precipitation Seasonality | Measure of variation in monthly precipitation | Positive values indicate more precipitation variability, negative values indicate less precipitation variability |
| Vegetation | |  |  |
|  | Tree Richness | Stand level measure of canopy tree richness |  |

Table S3. Dominant ant and spider species collected during the study. *Note: ant subfamilies are indicated in parentheses following species names.

| Species/Family* | | Proportion Individuals Collected |
| --- | --- | --- |
| Spiders | |  |
|  | *Eris militaris* (Salticidae) | 0.06 |
|  | *Theridion glaucescens* (Theridiidae) | 0.06 |
|  | *Anyphaena pectorosa* (Anyphaenidae) | 0.05 |
|  | *Spintharus flavidus* (Theridiidae) | 0.05 |
|  | *Euryopsis funebris* (Theridiidae) | 0.04 |
| Ants | |  |
|  | *Aphaenogaster mariae* (Myrmicinae) | 0.21 |
|  | *Camponotus nearcticus* (Formicinae) | 0.18 |
|  | *Lasius americanus* (Formicinae) | 0.11 |
|  | *Camponotus pennsylvanicus* (Formicinae) | 0.10 |
|  | *Prenolepis imparis* (Formicinae) | 0.10 |

Figure S2. Dendrogram of ant functional groups derived from the FDis function (FD package, R). Ant functional groups were delineated by setting the functional group number to 6, where we noted a clear break of the functional groups.

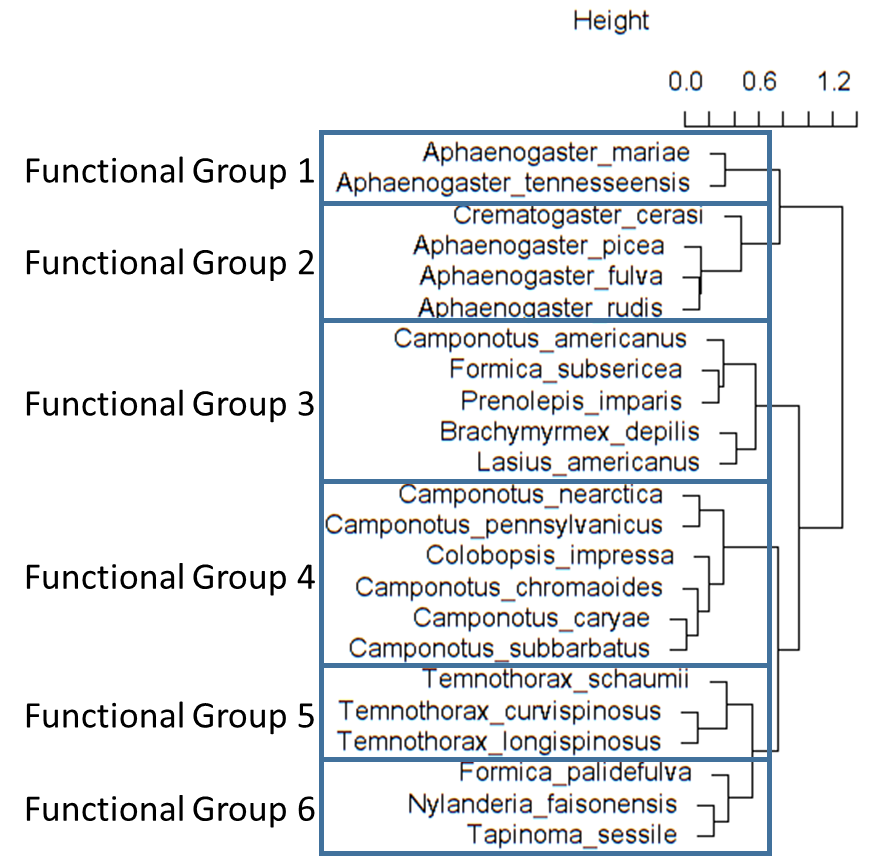

Figure S3. Presence-absence based rarefaction/extrapolation curves of (A) spiders and (B) ants collected across 96 trees. Solid lines are interpolated richness, points represent observed richness, while dashed lines are extrapolated richness. Curves were calculated with the iNEXT function and package (R). Shaded regions are 95% confidence intervals.

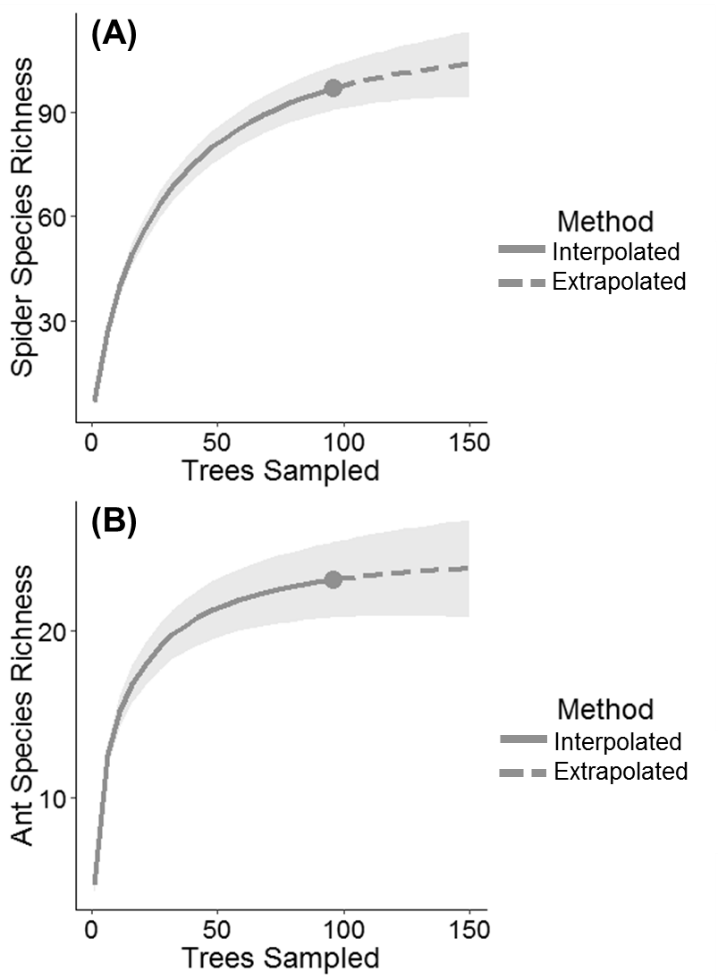
